# Effects of substrate availability on growth and metabolism in soil microbes: Insights from theoretical modeling of studies of the Warburg effect and substrate-induced respiration

**DOI:** 10.1101/2020.09.08.287813

**Authors:** Anshuman Swain, William F Fagan

**Affiliations:** Department of Biology, University of Maryland, College Park, MD 20742

**Keywords:** Substrate induced respiration, Warburg effect, Linear maximization model, Overflow metabolism, Carbon use efficiency

## Abstract

Carbon Use Efficiency (CUE) is a popular concept for measuring the efficiency of biomass production in different biological systems and, is frequently employed to understand effects of microbial processes on soil carbon dynamics. CUE in soil microbes is often measured through respiration-based studies, especially through the addition of a labile carbon substrate such as glucose. Therefore, exploring the response of microbial respiration to availability of labile substrates is crucial to understand microbial CUE in soils. In this work, we build upon a cellular model of the Warburg effect, where cells simultaneously utilize inefficient aerobic glycolysis/fermentation and efficient oxidative phosphorylation pathways for energy synthesis even at high oxygen availability, to predict microbial community response to various levels of substrate availability. We test our predictions systematically using a series of substrate-induced respiration (SIR) experiments to demonstrate prevalence of the Warburg effect in soil microbial communities. We further discuss the relevance of the underlying metabolic processes behind the Warburg effect in interpreting soil microbial CUE.

## 1. Introduction

Carbon Use Efficiency (CUE) of the soil microbial community, usually defined as carbon biomass produced per substrate carbon utilized, is considered an important variable that influences what proportion of organic carbon is released into the atmosphere as CO_2_ and what stays back as soil organic matter in (living or dead) microbial biomass (Billings and Ballantyne, 2013; Bradford, 2013; Hagerty et al., 2014). CUE depends strongly on the nature and quantity of available substrate and the compounds being produced, as well as on the physiology of the cells (e.g., active growth and division, survival when substrate availability is low, dormancy) (Dijkstra et al., 2015). However, CUE is usually considered to be low because of low carbon availability in soil and the supposedly recalcitrant nature of soil organic matter (Anderson and Domsch, 2010; Manzoni et al., 2012; Sinsabaugh et al., 2013; Reischke et al., 2015). Studies usually assume that the limited substrate available is used to satisfy basal metabolic demands for the cells with negligible growth. However, this assumption is not supported by numerous studies that find high values of CUE (Brant et al., 2006; Frey et al., 2013; Hagerty et al., 2014; Steinweg et al., 2008; Ziegler et al., 2005; Geyer et al., 2019), which sometimes approach pure culture values (Djikstra et al., 2015). Moreover, the assumption that soils are oligotrophic and that microbes are normally in a ‘starvation-survival lifestyle’ is flawed (Hobbie and Hobbie, 2013). Although most of the soil organic carbon (SOC) is dominated by recalcitrant carbon, the small portion involving labile carbon (or, low molecular weight carbon compounds) has very high turnover (Schneckenberger et al., 2008). Consequently, this labile carbon plays a major part in soil nutrient dynamics, and influences measurements of CUE.

CUE is often measured in soils using respiration studies that quantify the rate of mineralization of a substrate (mostly labile carbon compounds such as glucose) into CO_2_ or other gaseous or soluble simple compounds (Manzoni et al., 2012; Sinsabaugh et al., 2016). However, these respiration-based methods assume that CO_2_ emission in heterotrophic microbial communities is a good proxy for measuring microbial metabolic processes and biomass, and further assume that the proxy relationship is linear. Previous studies have reported the presence of non-linear effects of increased labile substrate concentration on CO_2_ emissions, especially the reduction of CO_2_ emissions at high substrate concentrations (Molenaar et al., 2009). This nonlinearity occurs because the rate of oxidative phosphorylation declines relative to other pathways as substrate concentrations increase. Eventually, at the highest substrate concentrations, aerobic glycolysis becomes the primary energy producing pathway, even in the presence of ample oxygen; this phenomenon is termed the Warburg effect (or overflow metabolism) (Schuster et al., 2015). This effect, initially discovered in cancer cells, has been well documented across a broad range of organisms (summarized in Swain and Fagan, 2018) including yeast (de Deken et al., 1966), bacteria (Basan et al., 2015, Zhuang et al., 2011, Molenaar et al., 2009), and various human cell types including activated lymphocytes, Kupffer cells, fast twitch muscle fibers, and microglial cells (Schuster et al., 2015, Vander Heiden et al., 2009). Here, we will focus on microbial (bacterial) respiration processes in the soil and its impact on CUE.

Although seemingly paradoxical at first, the Warburg effect occurs in bacterial cells due to three major trade-offs that occur between the two dominant energy producing pathways: oxidative phosphorylation (OP) and fermentation/aerobic glycolysis. These tradeoffs involve (1) the rate and yield tradeoff between OP and fermentation; (2) OP being a surface process whereas fermentation is volume-based; and (3) fermentation produces precursors to be used for growth whereas OP cannot do so directly (Molenaar et al., 2009; Zhuang et al., 2011; Schuster et al., 2015). Taking these three trade-offs into account, previous studies developed an array of frameworks including linear maximization models (Swain and Fagan, 2018) and flux balance analysis (Schuster et al., 2015) to explain the occurrence of Warburg effect at a cellular level.

Here, we use the cellular level linear maximization model from our earlier study (Swain and Fagan, 2018) to develop a set of hypotheses detailing the community-level consequences of the Warburg effect. We test these predictions using a series of simple substrate-induced respiration (SIR) experiments, and discuss these results in the context of CUE as a function of substrate availability in the soil and consequences for respiration-based studies.

## 2. Hypothesis Generation

Increased substrate availability tips trade-offs in favor of fermentative processes in bacterial cells at the expense of OP. In contrast, OP is favored at low substrate concentrations where efficiency of energy production is necessary to meet basal energy requirements (Swain and Fagan, 2018; Kempes at al., 2017). OP yields about 30 moles of ATP per mole of glucose consumed and is much more efficient compared to fermentation, which produces 2-4 moles of ATP per mole of glucose. However, fermentation is about two orders of magnitude faster than OP (Voet, 2004). Therefore, at high labile substrate availability, kinetic factors and availability of enzymatic centers would lead to an increased rate of fermentation. The enzymes that help facilitate OP, cytochrome oxidases, are located exclusively as transmembrane proteins, whereas enzymes that facilitate fermentation occur in the cytoplasmic space (Zhuang et al., 2011). To maintain membrane integrity, only a small proportion of the transmembrane space can be occupied by proteins; this limits the upper rate of OP to be much lower than fermentation. Moreover, at high substrate concentrations, transporters compete for membrane space with OP (Zhuang et al., 2011), thereby reducing OP. In addition, the products of fermentation facilitate biomass production (Vander Heiden et al., 2009) and therefore promote growth. However, the energy produced by fermentation is insufficient to support biosynthesis of all the carbon precursors produced through its enzymes, and energy produced from OP would be utilized in biosynthesis. Therefore, with increasing substrate concentrations, the total energy produced from both OP and fermentation would not suffice to process the fermentation-based products for biosynthesis. As a result, these excess products are expelled from the cell as overflow products, leading to the term overflow metabolism (or, Warburg effect) (Schuster et al., 2015).

These processes happen at a cellular level, but the surface-area-to-volume-ratio (henceforth termed Z) varies widely among individual cells, and therefore plays a major role in the prevalence of overflow metabolism as well as in relative growth, competitive substrate uptake, and minimum energy requirements (Swain and Fagan, 2018). Higher values of Z enable growth at lower substrate concentrations due to higher rate of substrate uptake at a given concentration. However, cells with lower Z values grow slower than those with larger Z values (Young, 2006; Swain and Fagan, 2018).

Given great diversity and high abundance of bacterial cells in soils, we can assume that Z is normally distributed in a sample of soil. This assumption takes into account both the size and shape of the cell, and controls the extent of overflow metabolism in each individual. We use the framework from a previous linear maximization model (see Swain and Fagan, 2018 for details), and apply it to this distribution of hypothetical cells which differ in Z (but, for simplicity, have the same volume) to calculate the CO_2_ mineralization rate from a simple substrate such as glucose. We assume the parameter values associated with various biochemical processes (see Swain and Fagan, 2018 for details) in the model to be normally distributed and randomly assigned.

For 100 starting substrate concentrations, we ran simulations of the model outlined by Swain and Fagan (2019) for 1 million different initializations each, using Latin hypercube sampling (LHS) (via the *lhs* package (Carnell, 2020) in R), to evaluate the effect of substrate availability on carbon usage in bacterial communities. Each simulation started with a set of parameter values for various cellular processes and an initial value of substrate concentration. The substrate was assumed to be homogeneously distributed and was depleted when consumed. Although the exact values of cumulative mineralization of carbon substrate to CO_2_ after a fixed time obtained using the Warburg effect model differed among simulations, they had the same underlying qualitative behavior (see Figure 1A).

**Figure 1:**
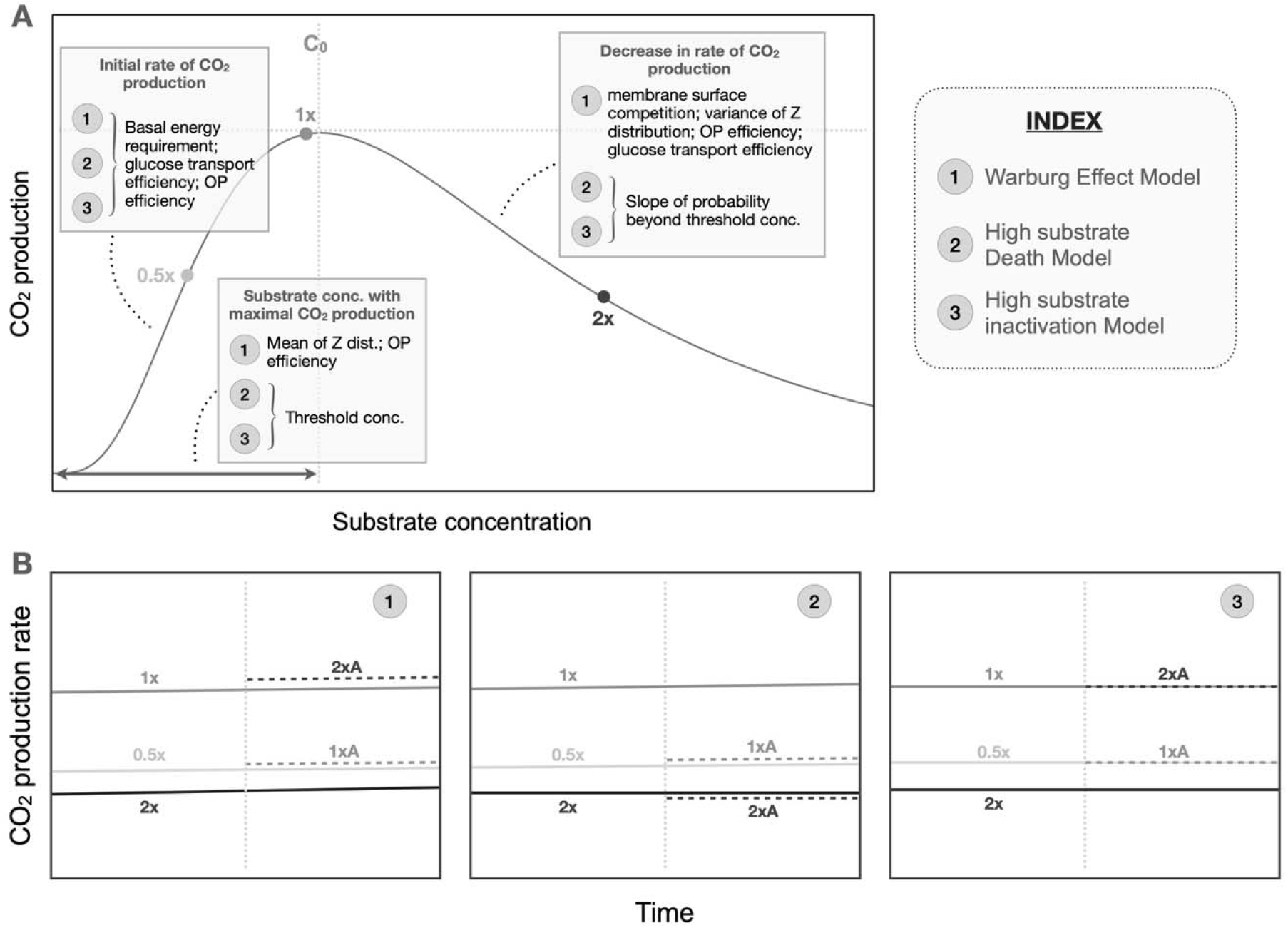
Theoretical predictions from our model: (1) CO_2_ production at different substrate concentrations and the factors that affect them (estimated using PRCC; see Fig. S1-S3) for three models discussed in the text; the values for cumulative CO_2_ production scaled with increasing time, as expected. The concentration where maximal CO_2_ (C_0_) production occurs is represented with a vertical dotted line. (2) Variations in CO_2_ production rates among the three models for three concentrations of substrate 0.5x, 1x and 2x (depicted using solid lines) followed by an abrupt halving of substrate concentration in 1x and 2x samples resulting in 1xA (1x altered) and 2xA (2x altered) samples (CO_2_ production rates are depicted in dashed lines).

We observed that the cumulative CO_2_ production in our Warburg Effect model increased smoothly with increasing substrate concentration until a certain concentration (let us call it - C_0_) where the maximum CO_2_ production occurred and, after which the production declined slowly (Fig 1A). We used a partial rank correlation coefficient (PRCC) via the R package *sensitivity*(Iooss et al., 2020) to identify which parameters affected different parts of the curve (Fig S1). We found that the initial rate of CO_2_ production increase depended negatively on the basal energy requirement, glucose transport efficiency, and OP efficiency. The substrate concentration where the maxima occurred depended on the mean value of the starting Z distribution, and OP efficiency. The rate of decrease of the curve beyond the threshold depended on the competition for membrane surface, variance of the starting Z distribution, OP efficiency, and glucose transport efficiency.

This behavior, seen in Fig. 1A, is also possible in two possible alternative scenarios: (1) microbes die due to very high substrate concentrations or, (2) microbes become inactive due for the same reason. Some previous studies have shown that high substrate concentration can result in cell inactivation or even death (Edwards, 1970; Han and Levenspiel, 1988). To simulate this, we built two simple models with similar resource usage patterns and cell distributions (and their properties) as the original Warburg effect model – one for cell death and one for cell inactivation. We assigned a threshold substrate concentration (referred to as the threshold parameter) at which the probability of death or becoming inactive (but alive) was 0.5 for the microbial population and increased linearly from zero substrate and, this slope of increasing probability is the second parameter that can be controlled in these two alternative models. We also removed the constraints associated with the Warburg effect (e.g., competition for membrane space, and upper limit of OP enzymes on the membrane) in these two models – so that the bacteria could make maximal use of the substrate input, but were subject to death or inactivation due to increased substrate levels.

We outline the parameters that affect the CO_2_ production in figure 1A (and provide details in Figures S1-S3). We can differentiate between the scenarios by abruptly changing our substrate regime where the value of CO_2_ production is greater than the maximal CO_2_ production rate in Figure 1A to one where it is lower. To systematically compare this, we selected a specific concentration (1x) such that it is slightly less than the concentration where maximal CO_2_ production occurs. Twice this concentration (2x) is then substantially greater than the concentration leading to maximal CO_2_ production, and 0.5x is far below that concentration (see Figure 1A). On abruptly changing the concentration from 1x to 0.5x and 2x to 1x in a given simulation, we see different responses in the three models (the Warburg Model, the Death model, and the Inactivation model; see Figure 1B). For example, because the biomass production rate is higher at higher substrate concentrations in the Warburg model, we observe altered 2x (2xA) and altered 1x (1xA) having higher respiration than that of 1x and 0.5x respectively. In the cell death model, we see higher respiration in altered concentrations, but only for 1xA and not for 2xA. This occurred because the reduction in respiration was actually due to cell death at 2x despite higher biomass production with increased substrate. The inactivation model showed either the same or lower values of respiration at 1xA and 2xA as compared to 0.5x and 1x respectively.

To validate our predictions, we used a system of simple substrate-induced respiration (SIR) experiments (Anderson and Domsch, 1973) to investigate two things. First, we examined the effect of substrate concentration on CO_2_ emission (after a fixed time) to look for the humped pattern predicted by the simulations. Second, we inspected the rate of CO_2_ production at three focal concentrations (as described above and in Figure 1B) to understand the humped patterns. Although more advanced kinetic respiration methods (such as Panikov and Sizova, 1996) are available, we used basic SIR as we expected it to be adequate for our purposes and easier to implement without any advanced equipment.

## 3. Materials and Methods

### 3.1 Sample collection and treatment

Soil samples used in this study were collected from three different sites each in North Bhubaneswar and adjacent Barang region in the state of Odisha, India. All six sites were non-agricultural and formed a part of a semi-forested landscape. The details about the soils in each plot appear in table 1. Soil cores were taken using a metal boring sampler (4 cm diameter) from five randomly chosen spots at each site. The soil from 1-8 cm below ground level was collected.

**Table 1:** Details of the soil at six sampling locations

| Property | Soil A | Soil B | Soil C | Soil D | Soil E | Soil F |
| --- | --- | --- | --- | --- | --- | --- |
| Collection Location | 20.397° N,<br>85.825° E | 20.399° N,<br>85.819° E | 20.400° N,<br>85.816° E | 20.366° N,<br>85.824° E | 20.367° N,<br>85.822° E | 20.370° N,<br>85.835° E |
| Clay % | 31 | 25 | 43 | 23 | 24 | 50 |
| Sand % | 35 | 33 | 31 | 38 | 47 | 29 |
| Silt % | 34 | 42 | 26 | 39 | 29 | 21 |
| pH<sub>1:5</sub> | 5.6 | 5.6 | 5.7 | 5.4 | 5.2 | 5.6 |
| Max. Water Holding Capacity (g kg<sup>-1</sup>) | 390 | 340 | 410 | 380 | 300 | 430 |
| Total Organic Carbon (g kg<sup>-1</sup>) | 35 | 29 | 32 | 28 | 24 | 30 |
| Total Nitrogen (g kg<sup>-1</sup>) | 2.7 | 2.3 | 2.5 | 2.3 | 1.8 | 2.9 |
| Total Phosphorus (g kg<sup>-1</sup>) | 0.5 | 0.5 | 0.4 | 0.4 | 0.3 | 0.4 |

The soils were passed through a 2 mm mesh to remove plant matter, roots, nematodes and other undesired material, and to homogenize the distribution of substrate and produce a more dispersed release of carbon dioxide. Soil properties (pH, water holding capacity, texture, total organic Carbon, total Nitrogen and total Phosphorus) were estimated, after open-air drying of the soil for 24 hours at 35° C, and samples were then stored at 5° C in aerated plastic bags. Soil water-holding capacity was measured at matric potential 10 kPa following previous studies (Wilke, 2005). A 1:5 soil/water suspension ratio was used to estimate soil pH after 60 minutes shaking at 22 °C. Total organic Carbon was measured using the Walkley-Black procedure and multiplied with a factor of 1.33 to adjust for the carbon recovery rate (Schumacher, 2002). Total Nitrogen was estimated using the Kjeldahl digestion method (Taylor, 2000), and Total Phosphorus was estimated using the fusion method (Olsen and Sommers, 1982).

### 3.2 Substrate Induced Respiration and test for Warburg effect

All samples were kept at 22° C for 24 hours before assay and were subjected to three sets of trials. For each of the thirty soil samples (five for each of the six sites), the following basic protocol was followed across all trials: 10g of sieved soil was put in a 25 mL dry Erlenmeyer conical flask, with sufficient headspace (15-20 mL) to ensure there was no oxygen limitation. Glucose solution of a desired concentration was added with frequent mixing, keeping total (deionized) water added to the sample at 50 percent of the maximum water holding capacity. The mouth of the flask was corked using a rubber structure with a tube inserted through it and the other end of the tube ended in a corked conical flask containing 30 mL 0.01 M KOH solution. The setup was incubated at 22° C. The KOH solution, after a stipulated time as needed for a specific set of trials, was treated with 0.03 M Barium nitrate solution and titrated against 0.005 M HCl solution to estimate CO_2_ emission. Three replicates of every trial were performed for each of the 30 soil samples.

For the first set of trials, the basic protocol was performed for differing concentrations of glucose ranging from 0.05 g to 0.80 g per 10 g of soil (with increments of 0.05 g of glucose between samples). The cumulative total CO_2_ emission was measured after 4, 5 and 6 hours (with 3 replicates for each soil for each time point). We also maintained two control groups (with three replicates each for each of the three end timepoints), one with the addition of water to the normal soil, and the other to autoclaved soil at 0.5 maximum water holding capacity. Final estimates of respiration were calculated by subtracting the respiration from the normal soil without glucose addition from the respective sample. Autoclaved soil samples were maintained to see if there was any abnormal respiration.

The second set of trials were exact replicates of the first set in two groups except for the addition of 1000 ppm chloramphenicol and 2000 ppm cycloheximide combined with 50 mg of talc into the soil (and mixed well) to function as selective inhibitors for bacterial and fungal growth, respectively (based on Nakamoto and Wakahara, 2004). Their respective controls were also amended with chloramphenicol and cycloheximide. Although these treatments are not completely effective (Nakamoto and Wakahara, 2004) and there might be streptomycin-resistant bacteria and cycloheximide-resistant fungi in soils, such colonies are usually slow growing and small (Anderson and Domsch, 1975). Consequently, these experimental treatments would give us a good estimate of bacterial and fungal biomass separately.

From the theoretical predictions (Fig 1A), we expected a dip in the production of carbon dioxide after a certain concentration C_0_ of glucose (see Hypothesis Generation section). Using the data from the first two sets of trials, we estimated this concentration C_0_ and further focused only on the bacterial respiration (i.e., cycloheximide treatment) for two major reasons. The first reason is that our models are based on bacterial metabolic mechanisms and extrapolating them to all soil organisms could be inappropriate. Second, anti-bacterial compounds like chloramphenicol and streptomycin can stimulate microbial respiration, especially fungal respiration, which can cancel the decline seen in bacterial respiration (Johnson et al., 1996; Lin and Brooks, 1999). Therefore, looking at the cycloheximide treatment in the third set of trials is the most appropriate, as it involves mostly bacterial respiration.

For doing the same, four specific glucose concentrations were taken for each soil sample, as described in the hypothesis generation section (Fig 1A-B): a base concentration (1x) was chosen such that it was less than C_0_ (the concentration of glucose where maximal CO_2_ release happens) but twice the concentration (2x) was greater than C_0_.

Processing data from first two sets of trials, we fixed 1x to be 0.30 g glucose/10 g of soil (see Fig 2A-F). For each soil sample, we established SIR experiments for glucose concentrations of 0.15 g (0.5x), 0.30 g (1x) and 0.60 g (2x) per 10 g of soil and performed the titration every 30 minutes starting at 0.5 hr timepoint and ending at 7 hr for three replicates of a given sample. After periods of 4, 5 and 6 hours, 10 g of autoclaved soil (with cycloheximide amendment and water added at 0.5 maximum water holding capacity) was added to one set of three replicates each of 1x and 2x series of samples. CO_2_ emission rates were observed at the same 30-minute observation intervals. The addition of autoclaved soil functions as a diluter, thus reducing the substrate concentration to half – therefore making the 2x runs into 1x and 1x into 0.5x. For every soil we had 42 (3 x 14 time-points), 78 (3 x 14 time-points + 3 x 12 amendment samples (6 for 4 hr, 4 for 5 hr and 2 for 6 hr)) and 78 replicates for 0.5x, 1x and 2x respectively. We also maintained 78 replicates of each of the two control groups (i.e. normal soil and autoclaved soil) with cycloheximide amendment. A similar treatment (addition of autoclaved soil) was done to 3 replicates each of the two control groups for comparison after 4, 5 and 6-hour timepoints. To estimate the respiration after addition of autoclaved soil to 1x and 2x concentrations, we subtracted the respective respiration from autoclaved soil-added normal soil sample control. To estimate the rate, we subtracted the total CO_2_ produced at a given time-step from that of the previous time-step.

**Figure 2:**
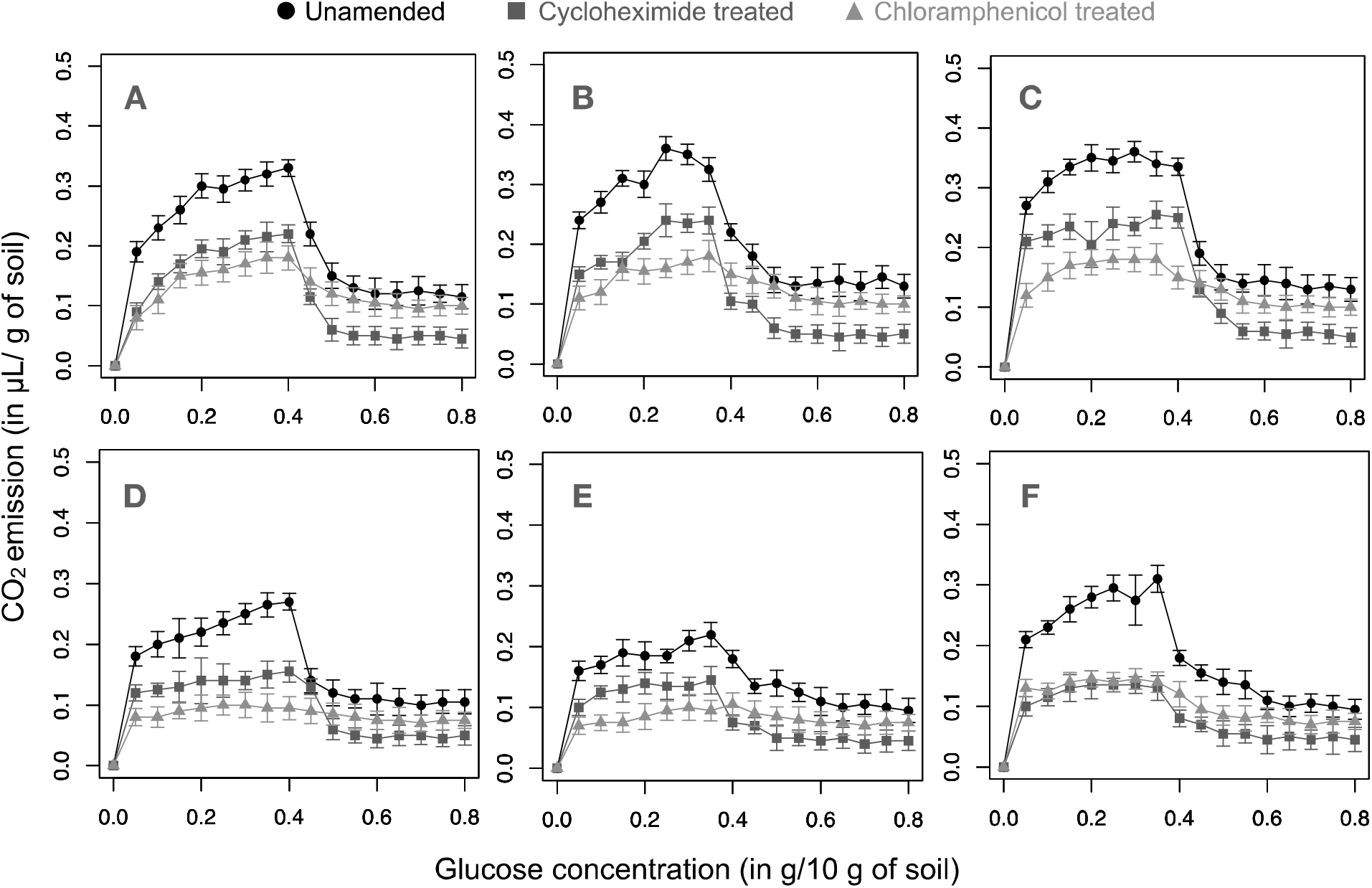
Cumulative SIR emissions from different soils A-F (representing soil of the same letter, respectively; see Table 1) after 4 hours of incubation at various concentrations of glucose and three different treatments for each soil – unamended soil (total biomass), cycloheximide treatment (primarily bacterial biomass) and chloramphenicol treatment (primarily fungal biomass). Note that the decreases in cycloheximide treated soils were larger and faster than their corresponding chloramphenicol treated samples.

All comparisons and analyses were performed using R statistical software. For comparing the emissions from the 0.5x, 1x and 2x series of data, we used paired t-tests to identify significant differences.

## 4. Results

Cumulative carbon dioxide emissions due to total, fungal, and bacterial soil microbial biomass after 4 hours of incubation are plotted in figure 2. Comparable emissions results after 5 and 6 hours of incubation appear in figures S4 and S5, respectively. As predicted by simulations (Fig. 1A), CO_2_ production from the same soil sample decreased beyond a threshold substrate concentration and saturated to a lower value at all the six soil sites and for all three treatments. The threshold concentration varied slightly across the six soil samples, which may be a function of the microbial composition and soil properties. These basic trends in our experimental results match predictions described by our model in Figure 1A, even though the model was specifically made for bacterial metabolism.

Respiration in the cycloheximide-treated soils (which involved predominantly bacterial respiration) underwent a sharper decrease than the respiration in the chloramphenicol-treated soils (which involved predominantly fungal respiration) (Fig. 2A-F). This pattern suggests that the decrease in overall microbial respiration was likely caused primarily by the decrease in predominantly bacterial respiration. The sum of the cycloheximide- and chloramphenicol-treated soil respirations together exceeded the total untreated/unamended microbial respiration (Fig. 2A-F).

The temporal carbon dioxide emissions from the third set of trials where autoclaved soil was added after 4 hours is shown in figure 3A-F. The results of soil addition after five and six hours show similar responses across all soils (Figs. S4, S5). The addition of the autoclaved soil caused the respiration level of the soil to increase for 2x concentration samples (2xA), matching and even exceeding the CO_2_ production levels of 1x concentration. The same addition to 1x concentration (1xA) samples reduced the CO_2_ levels to just more than the CO_2_ levels produced by the 0.5x concentration; this observation held true for all six sites. Paired t-tests (for all 4, 5 and 6 hr trials) showed that the emissions from the 2xA samples were significantly higher than 1x sample (p<0.05 for soils A, B, D and F for all the three timepoints; p<0.05 for C and E for 5 and 6 hr timepoints; p<0.1 for C and E for 4 hr timepoint) Likewise, emissions from 1xA samples were significantly higher than 0.5x samples (p<0.05 for soils B and F for all the three timepoints, and for soils A, C and E for 5 and 6 hr timepoints; p<0.1 for all other cases).

**Figure 3:**
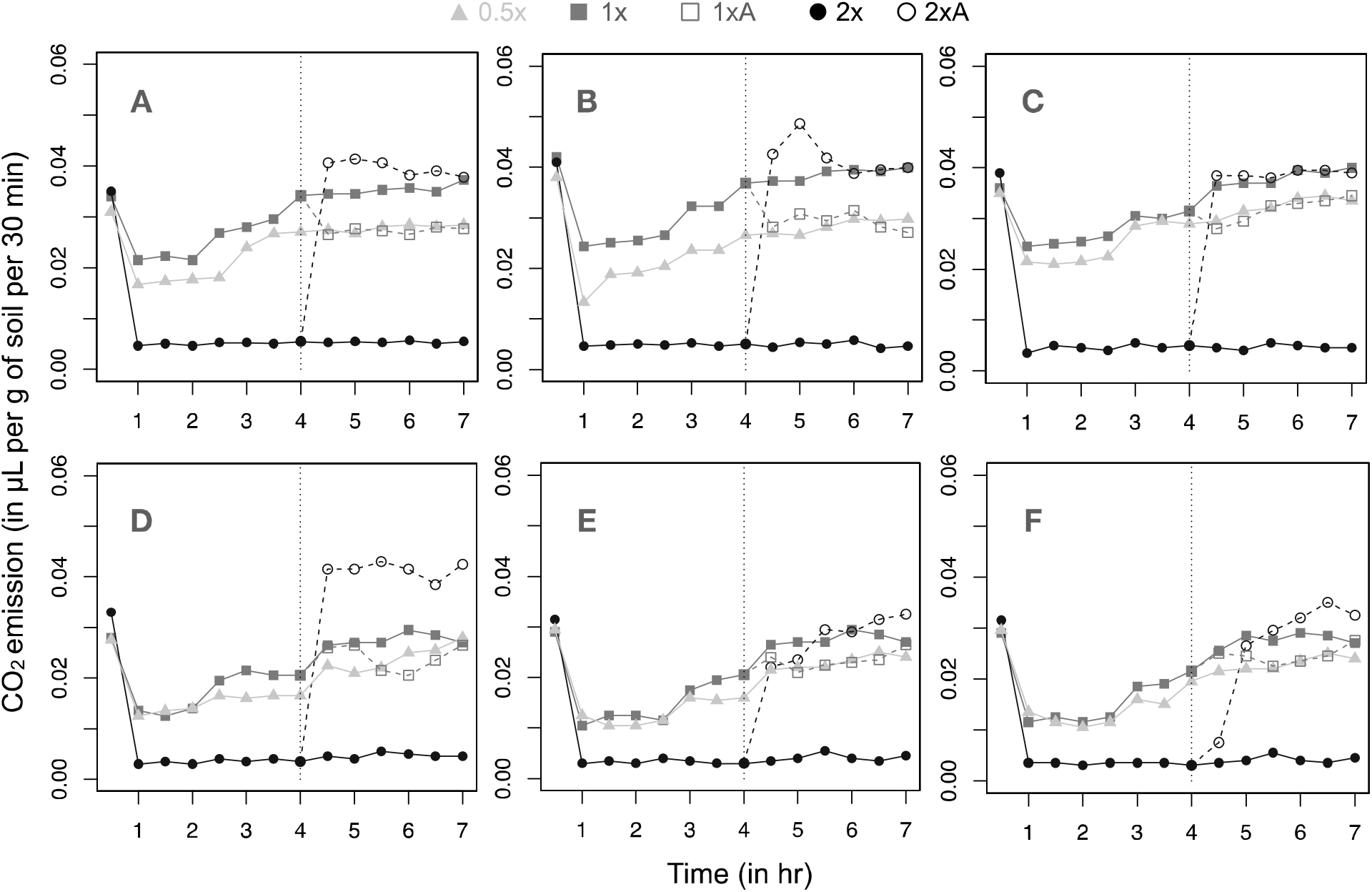
SIR emission rate values for cycloheximide-amended soil samples, with pre-determined concentrations of glucose: 0.5x (0.15g/10g of soil), 1x (0.3g/10g of soil) and 2x (0.6g/10g soil). Autoclaved soil was added after 4 hours (denoted by clack vertical line) in 1x and 2x samples, resulting in 1xA (1x altered) and 2xA (2x altered) samples respectively.

## 5. Discussion

The first two sets of trials for all accumulation periods (4, 5, and 6 hours of incubation) matched the qualitative shape of the curve predicted by our model (Fig 1A, Fig 2A-F, Fig S4A-F, Fig S5A-F). We observed that the additive total of CO_2_ emissions from the cycloheximide- and chloramphenicol-treated soils was greater than the total for the unamended soils (Fig 2A-F, Fig S4A-F, Fig S5A-F). This may be due to the presence of resistant microorganisms - thus contributing to both the measurements. Although our model is specifically made for bacterial metabolism, certain predictions from it might be applicable to fungal processes. However, for the sake of clarity, we focused on cycloheximide-amended soil (primarily bacterial biomass) for our third set of trials.

The third set of trials were conducted to differentiate among mechanisms that could have resulted in the curve observed in Fig 1A and the cumulative CO_2_ incubations (Fig 2A-F, Fig S4A-F, Fig S5A-F), allowing us to compare against the predictions for three possible models (the Warburg model, the cell death model, and the cell inactivation model) in Fig 1B. Based on the results (Fig 3A-F, Fig S6A-F, Fig S7A-F) for the third trials and the measured statistical differences between the two pairs (1x, 2xA) and (0.5x, 1xA) – the model that matched the data best was the Warburg model. A few cases were statistically inconclusive (i.e., p<0.1 but not <0.05), and we could not rule out the cell inactivation model completely.

It is also possible that different sets of microbes were active at different substrate regimes and that might cause small differences in respiration, as seen in previous studies (Derrien et al., 2014). Our model assumption in which the microbial community can be characterized as having a distribution of Z (surface area to volume ratio) values takes care of this scenario. By allowing microbes with different Z, we allow for a wide range of different energy requirements and growth regimes (Swain and Fagan, 2018). Although this normal distribution is a good approximation. Nevertheless, because our aim here was to create a framework for understanding the Warburg effect as a function of the responses of microbial processes to labile substrate availability, this represents a reasonable starting point. To capture a certain breadth of this complexity, we performed the experiments on six different soils with different physical and chemical properties (Table 1), and we observed similar trends of observations in soils across the six sampled sites. A further step in this direction might be to understand the consequences of the Warburg effect in different soil types and across different physical and chemical gradients.

We are not the first to see these effects in SIR-based studies. Indeed, past research has uncovered results similar to our first set of trials (e.g. Schneckenberger et al., 2008; Tian et al., 2015), where mineralization of glucose into CO_2_ follows a humped curve as a function of substrate concentration (Fig 1A). However, the questions in those studies were quite different from ours here, where we were primarily interested in understanding the implications of the Warburg effect (and its associated trade-offs) in the context of soil microbial community ecology. For example, our results can be assimilated in terms of carbon use efficiency (CUE), which is widely used in the soil literature (Lipson, 2015). Fig 1A can be repurposed to think about CUE as a function of substrate concentration by calculating the biomass C produced as a function of the input of cellular substrate C using the Warburg model (see Swain and Fagan, 2018; Fig 4).

**Figure 4:**
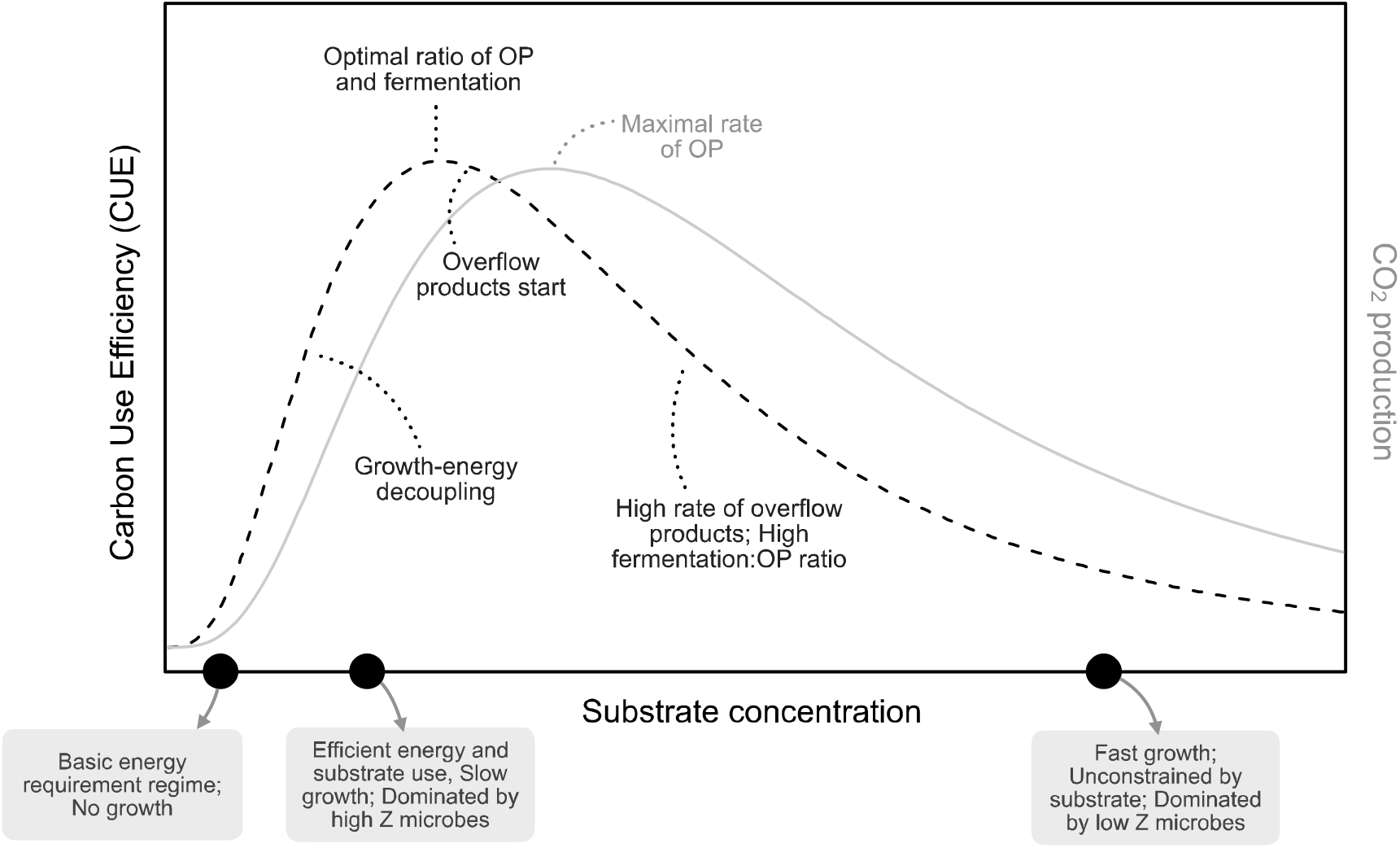
Conceptual model of Carbon Use Efficiency (CUE) as a function of substrate concentration

The overall trend of CUE represented here is similar to the model presented in a previous work by Lipson (2015), which was based on a different population-level tradeoff between metabolic rate and growth (Lipson, 2015; Beardmore et al., 2011) (Fig 4). At low substrate concentrations, microbial metabolism is dominated by high Z microbes, which have higher substrate use efficiency but slow growth. As the substrate availability increases, more cells cross the energy requirement barrier to achieve growth with high CUE. This continues until a certain threshold concentration of substrate is reached, at which fermentation-induced biomass production utilizes all precursors based on total energy availability. Beyond this threshold, the total energy produced is not sufficient to use all the precursors formed in glycolytic pathways – leading to overflow products. Even though the maximal OP rate might be reached at a higher substrate value, it is the ratio of fermentation-to-OP that matters. The higher the ratio, the larger the amount of overflow products that is expelled from the cell, leading to low CUE.

This trend in CUE can lead to different ecological ‘strategies’ among the members of microbial communities – both spatially and temporally. For example, inefficient fast-growers, efficient slow-growers and extremely efficient very-slow-growers could all coexist with each group predominating for particular levels of environmental substrate availability. Many previous studies also show that high CUE is representative of cooperative populations, and these can be particularly successful in spatially structured environments with varying resource regimes (Pfeiffer et al., 2001; Kreft and Bonhoeffer, 2005). However, the tradeoff between rate and yield can lead to coexistence of different microbial growth strategies even in a well-mixed environment (Beardmore et al., 2011). Temporal variations in substrate availability can also dynamically alter the microbial composition such as increased slow growing population at lower substrate concentrations (which happen with increased time of incubation) (see Derrien et al., 2014). The value of CUE also hinges on limiting nutrients such as N and P (Sinsabaugh et al., 2016; Sterner and Elser, 2002). In this work we have not considered the limitation imposed by N and P availability in the soils above the substrate availability; this topic remains open for further exploration.

Deciphering the relationships among the Warburg effect, CUE, and soil microbial respiration can be crucial for several reasons. For example, numerous methods that are commonly used to calculate soil microbial biomass and substrate utilization models assume merely a linear response of respiration to changes in substrate concentration (Schnecker et al., 2019). If this relationship is commonly nonlinear, as evident in scenarios where the Warburg effect is active, the applications employing the respiration-substrate concentration relationship could be over- or under-estimating aspects of microbial dynamics that can have downstream implications for the environment or the study system. In other words, understanding the nonlinear response of microbial respiration to changes in substrate concentration in an experimental setting can improve analyses that use substrate priming techniques for biomass estimation (Dilly and Zyakun, 2008) as well as understanding differential microbial community dynamics (Derrien et al., 2014). In addition, exploring these non-linear relationships can help explain the effects of sudden/periodic substrate/nutrient addition to soil systems such as leaf litter addition during fall season in temperate soils, rewetting of arid soils (allowing for pulsed usage of substrates), or other forms of nutrient periodicity and variability (Carrero-Colon et al., 2006; Lennon and Cottingham, 2008).

## Acknowledgements

We would like to thank Jake Weissman, and Margaret Palmer for their discussions about the manuscript. This research did not receive any specific grant from funding agencies in the public, commercial, or not-for-profit sectors. No conflict of interest to be declared.

## Supplementary Figures

**Figure S1:**
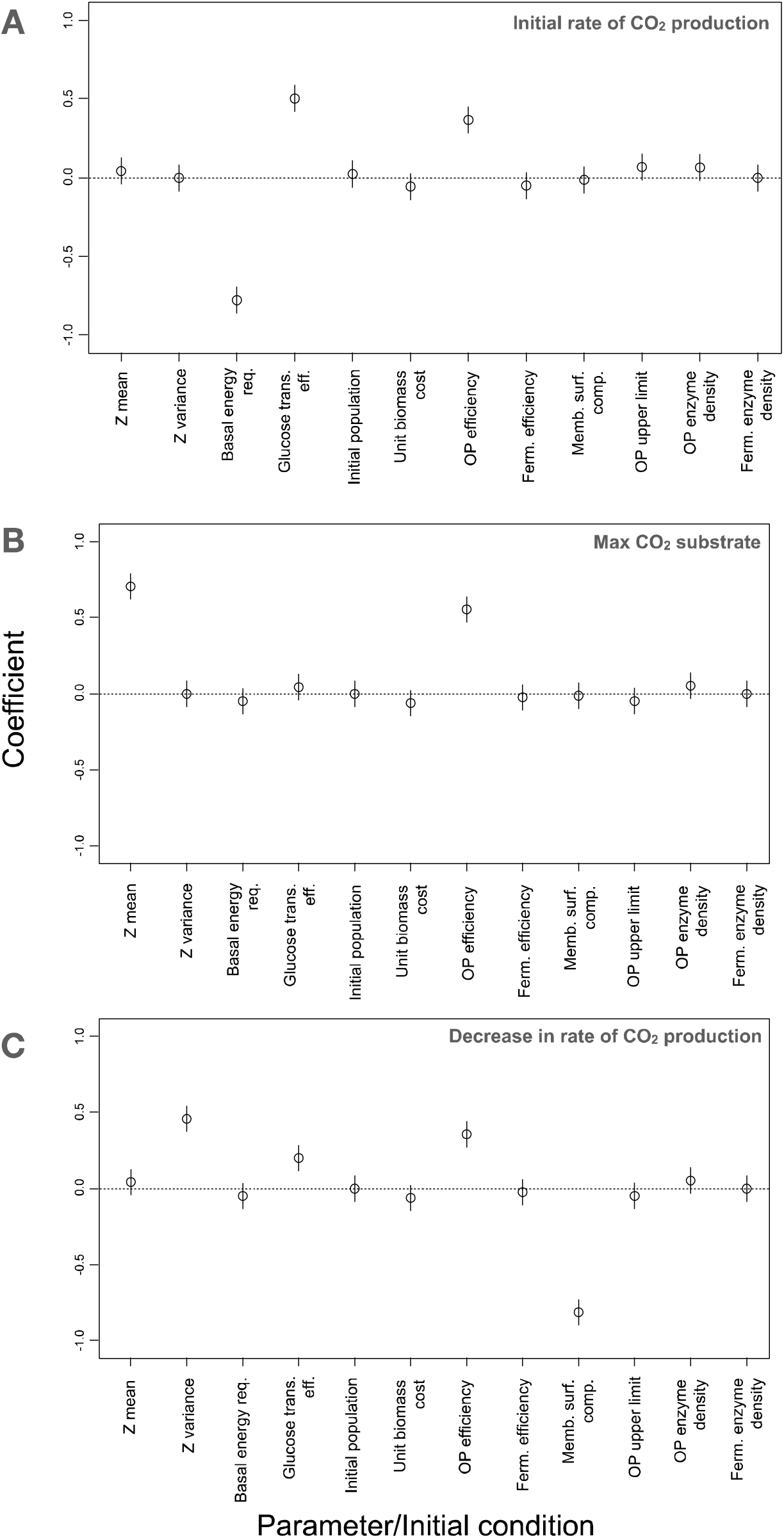
PRCC coefficients for the Warburg model for three properties of CO_2_ production

**Figure S2:**
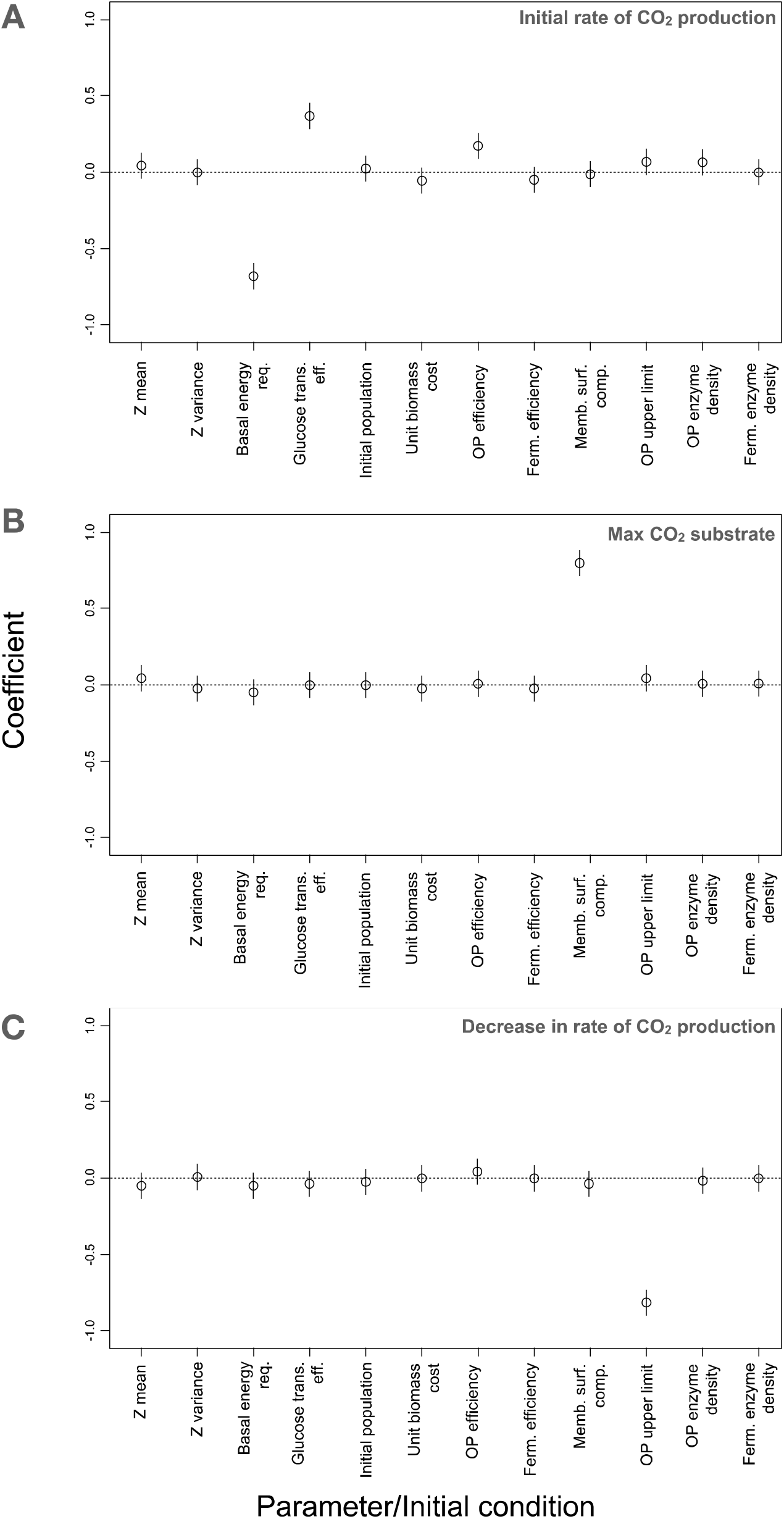
PRCC coefficients for the Death model for three properties of CO_2_ production

**Figure S3:**
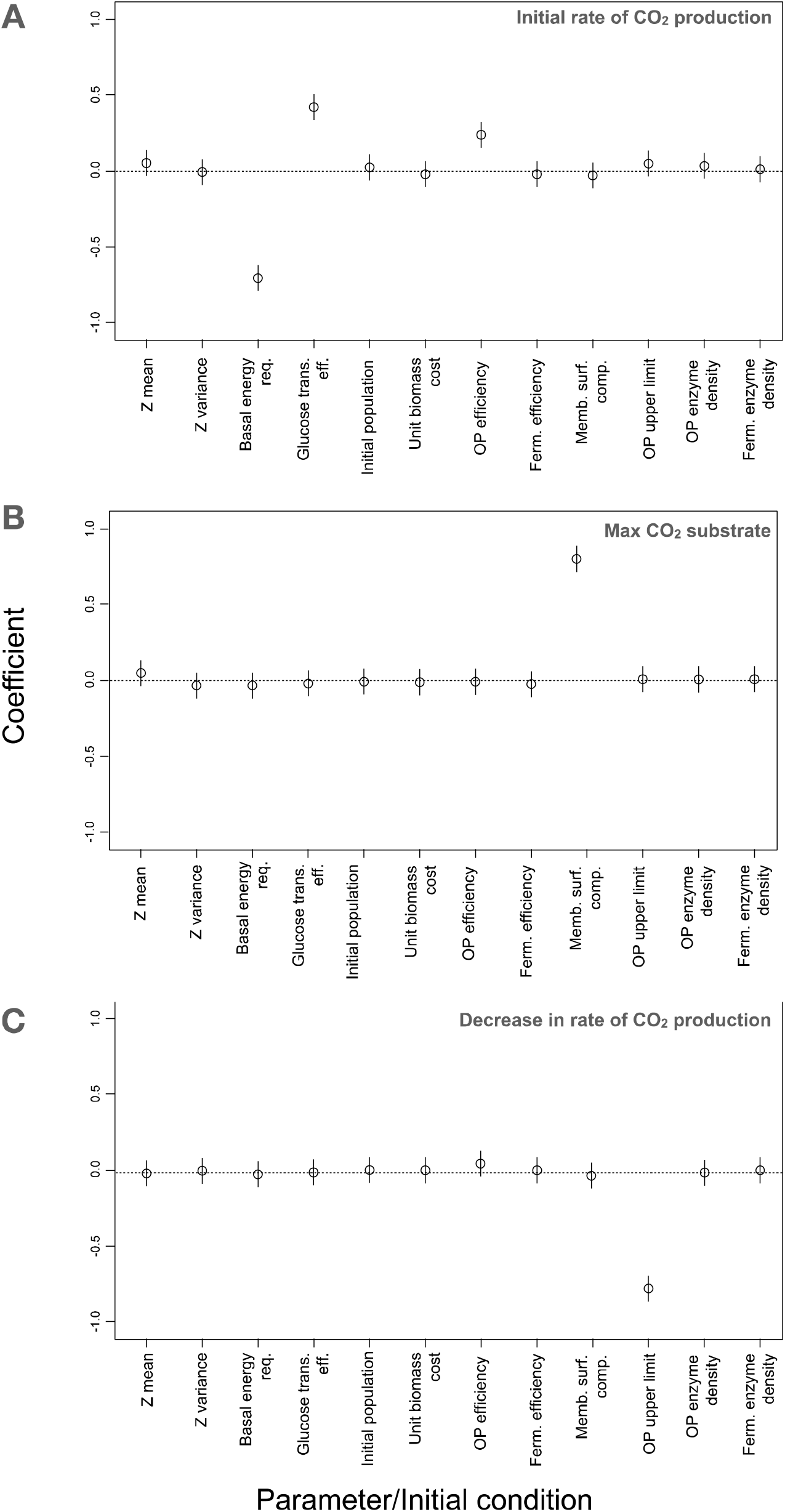
PRCC coefficients for the Inactivation model for three properties of CO_2_ production

**Figure S4:**
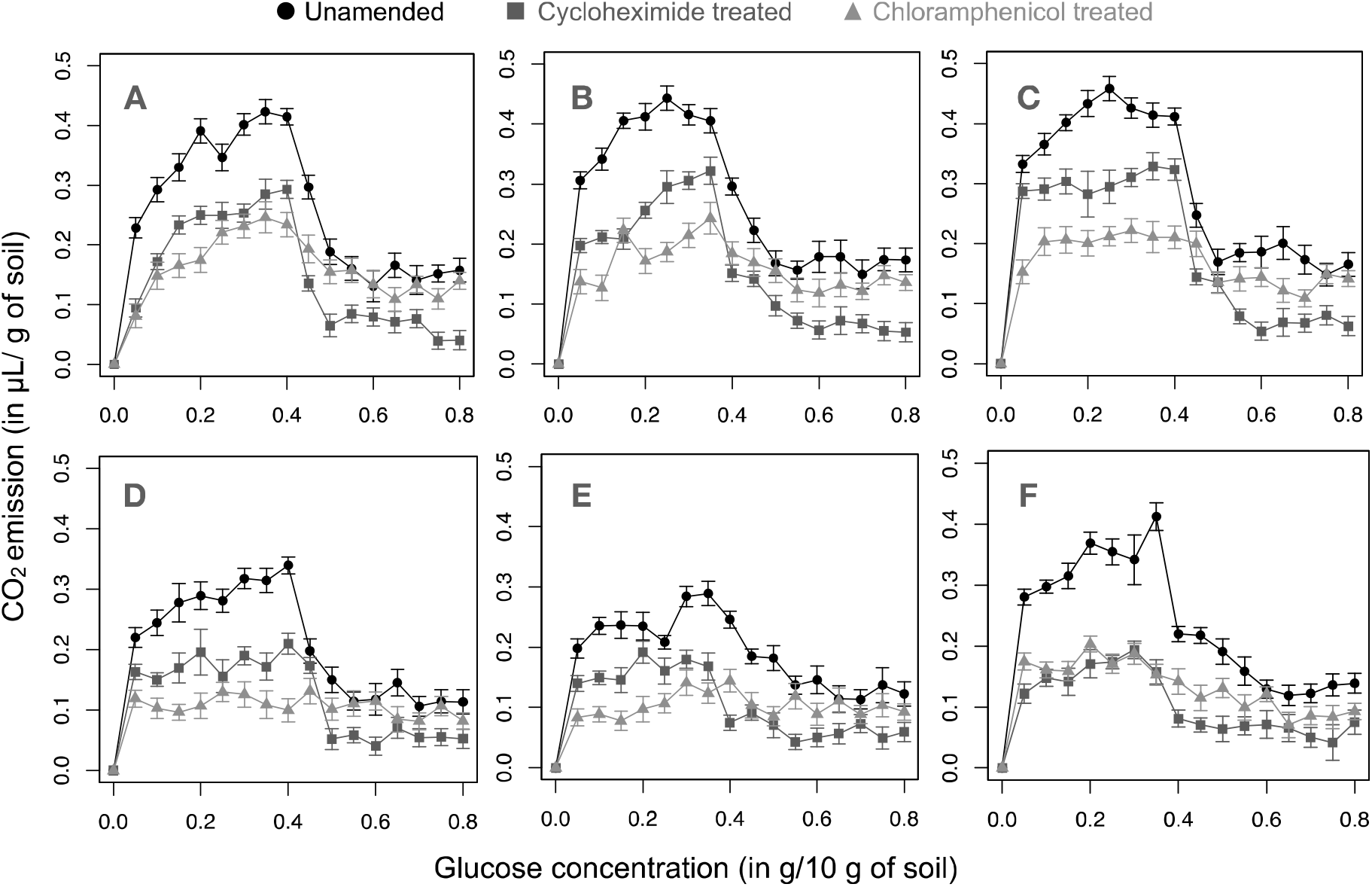
Cumulative SIR emissions from different soils A-F (representing soil of the same letter, respectively; see Table 1) after 5 hours of incubation at various concentrations of glucose and three different treatments for each soil – unamended soil (total biomass), cycloheximide treatment (primarily bacterial biomass) and chloramphenicol treatment (primarily fungal biomass).

**Figure S5:**
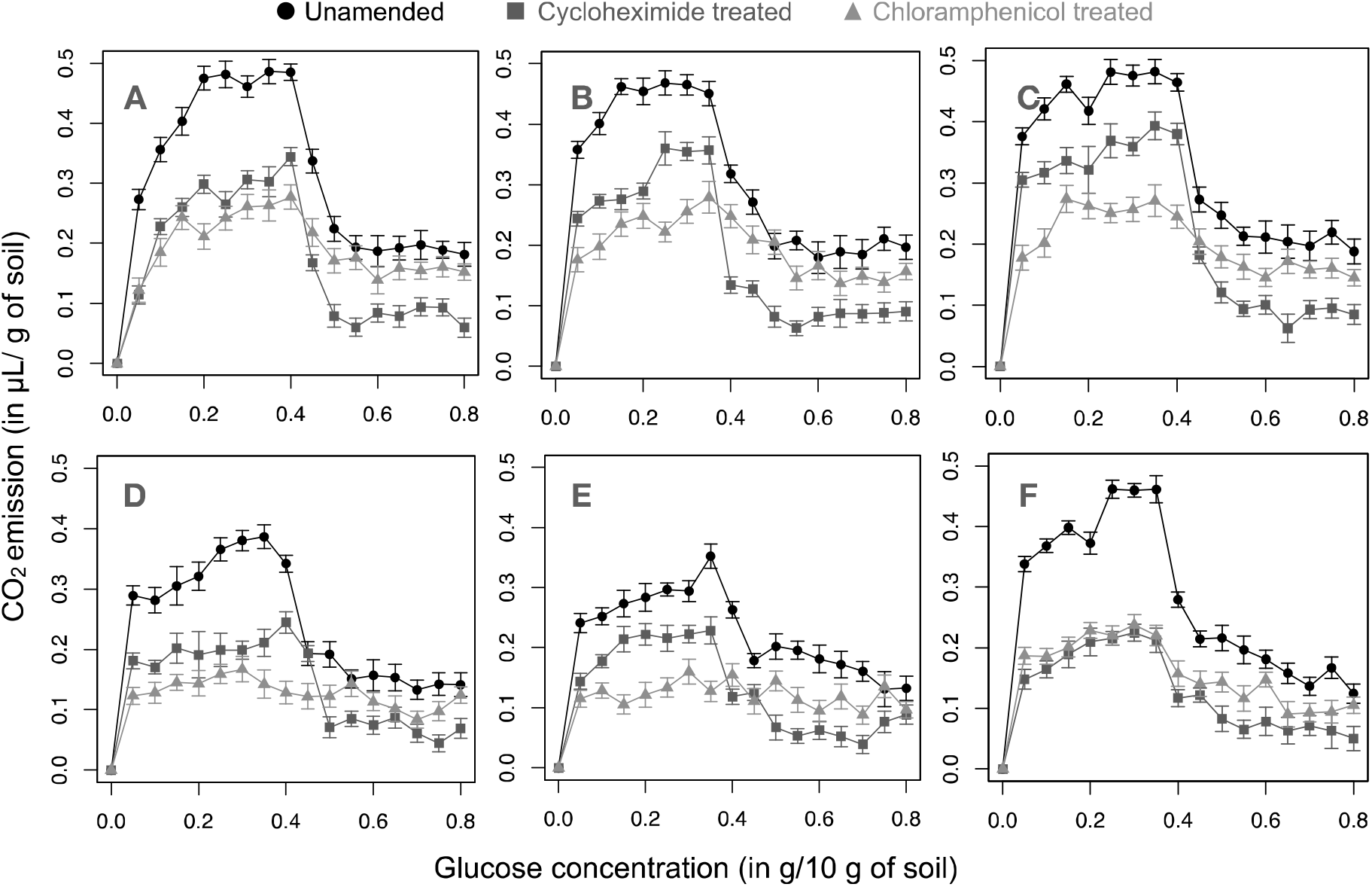
Cumulative SIR emissions from different soils A-F (representing soil of the same letter, respectively; see Table 1) after 6 hours of incubation at various concentrations of glucose and three different treatments for each soil - unamended soil (total biomass), cycloheximide treatment (primarily bacterial biomass) and chloramphenicol treatment (primarily fungal biomass).

**Figure S6:**
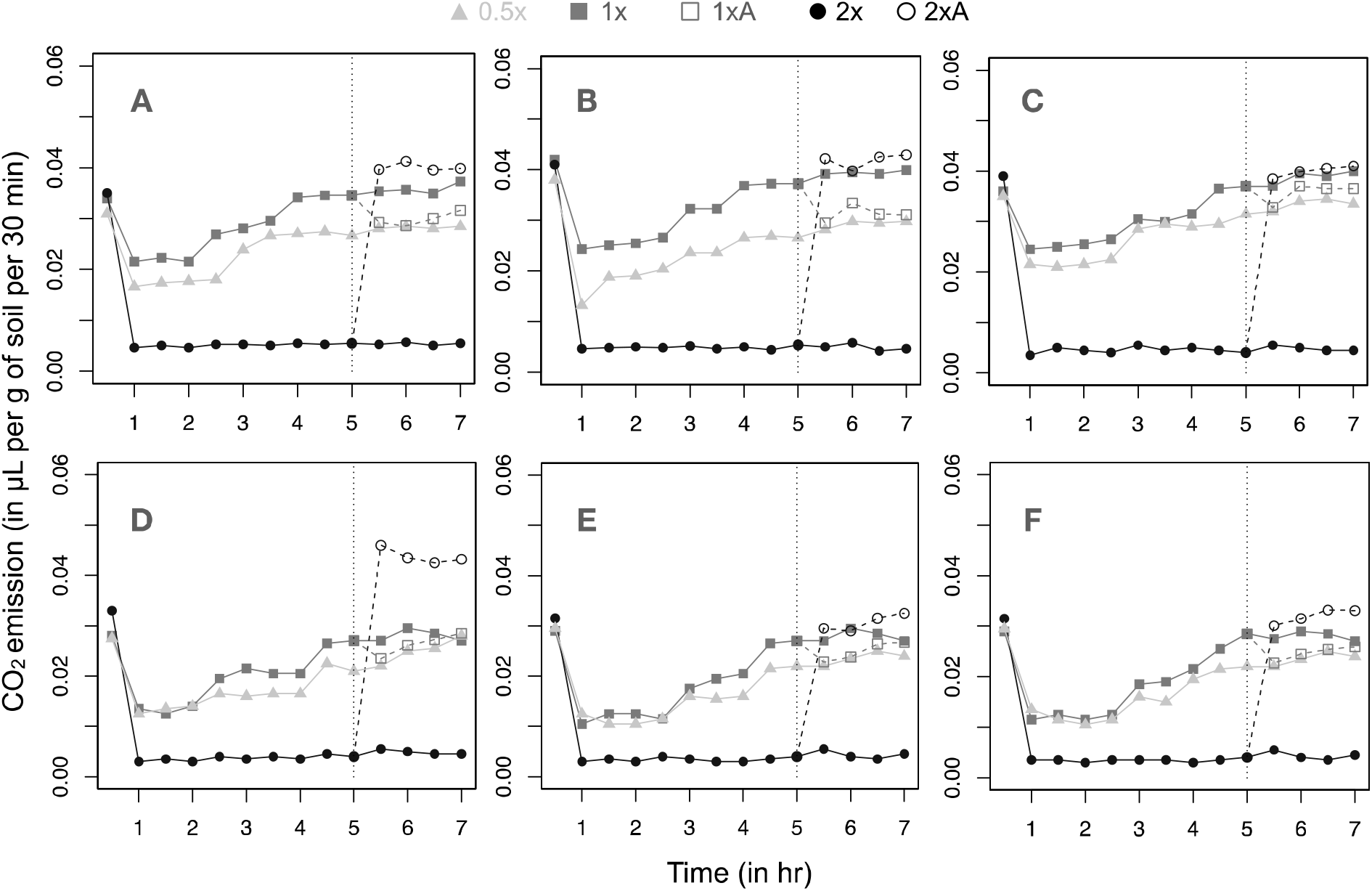
SIR emission rate values for cycloheximide amended samples of soils, with pre-determined concentrations of glucose: 0.5x (0.15g/10g of soil), 1x (0.3g/10g of soil) and 2x (0.6g/10g soil). Autoclaved soil was added after 5 hours (denoted by clack vertical line) in 1x and 2x samples, resulting in 1xA (1x altered) and 2xA (2x altered) samples respectively.

**Figure S7:**
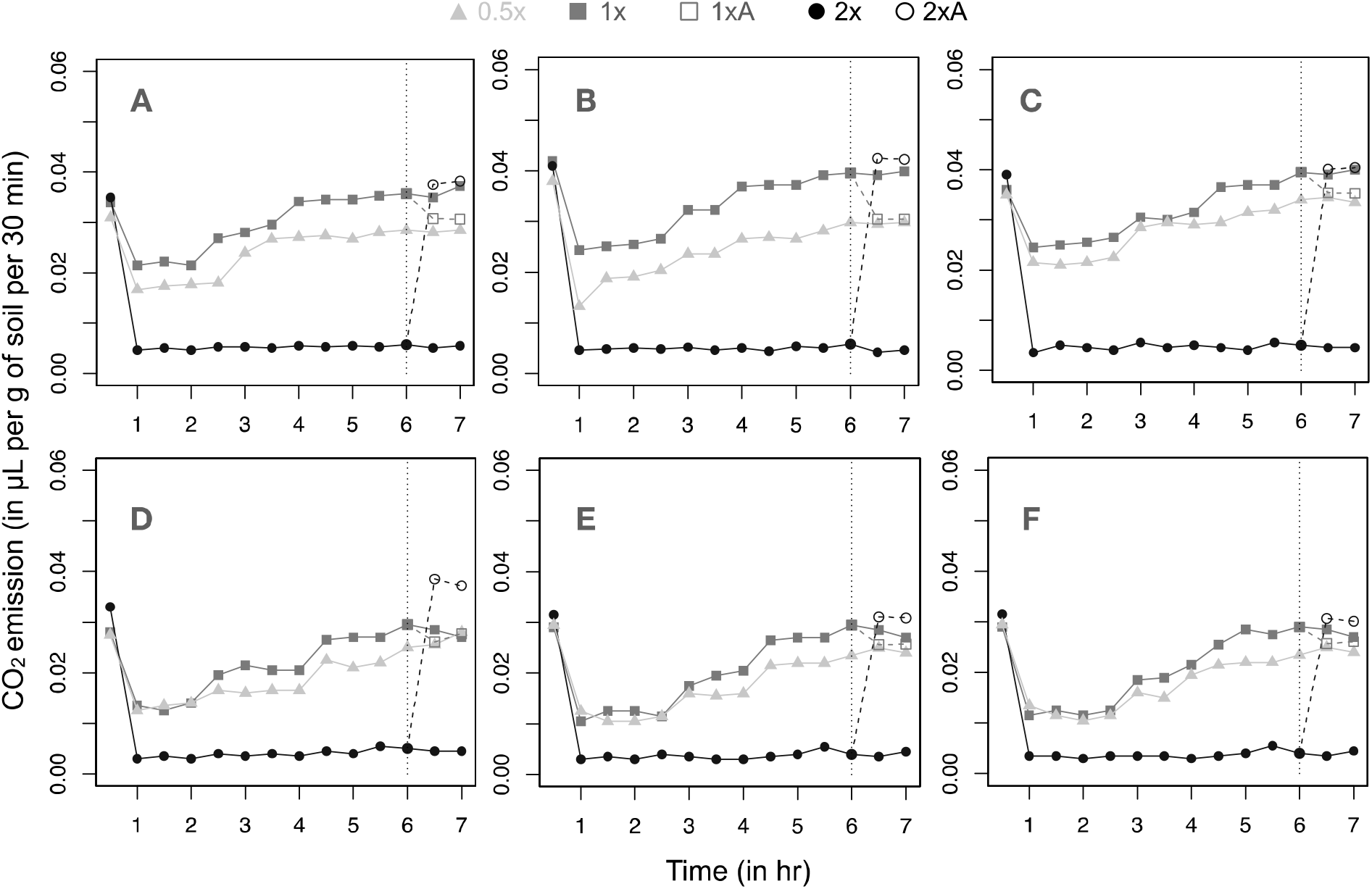
SIR emission rate values for cycloheximide amended samples of soils, with pre-determined concentrations of glucose: 0.5x (0.15g/10g of soil), 1x (0.3g/10g of soil) and 2x (0.6g/10g soil). Autoclaved soil was added after 4 hours (denoted by clack vertical line) in 1x and 2x samples, resulting in 1xA (1x altered) and 2xA (2x altered) samples respectively.

## Notes

### Competing Interest Statement

The authors have declared no competing interest.

## References

1. Anderson, J.P.E. and Domsch, K.H., 1975. Measurement of bacterial and fungal contributions to respiration of selected agricultural and forest soils. Canadian Journal of Microbiology, 21(3), pp.314–322.

2. Anderson, T.H. and Domsch, K.H., 2010. Soil microbial biomass: the eco-physiological approach. Soil Biology and Biochemistry, 42(12), pp.2039–2043.

3. Basan, M., Hui, S., Okano, H., Zhang, Z., Shen, Y., Williamson, J.R. and Hwa, T., 2015. Overflow metabolism in Escherichia coli results from efficient proteome allocation. Nature, 528(7580), pp.99–104.

4. Beardmore, R.E., Gudelj, I., Lipson, D.A. and Hurst, L.D., 2011. Metabolic trade-offs and the maintenance of the fittest and the flattest. Nature, 472(7343), pp.342–346.

5. Billings, S.A. and Ballantyne IV, F., 2013. How interactions between microbial resource demands, soil organic matter stoichiometry, and substrate reactivity determine the direction and magnitude of soil respiratory responses to warming. Global Change Biology, 19(1), pp.90–102.

6. Bradford, M.A., Keiser, A.D., Davies, C.A., Mersmann, C.A. and Strickland, M.S., 2013. Empirical evidence that soil carbon formation from plant inputs is positively related to microbial growth. Biogeochemistry, 113(1-3), pp.271–281.

7. Brant, J.B., Sulzman, E.W. and Myrold, D.D., 2006. Microbial community utilization of added carbon substrates in response to long-term carbon input manipulation. Soil Biology and Biochemistry, 38(8), pp.2219–2232.

8. Carnell, R., 2020. lhs: Latin Hypercube Samples. R package version 1.0.2. https://CRAN.R-project.org/package=lhs

9. Carrero-Colón, M., Nakatsu, C.H. and Konopka, A., 2006. Effect of nutrient periodicity on microbial community dynamics. Applied and environmental microbiology, 72(5), pp.3175–3183.

10. De Deken, R.H., 1966. The Crabtree effect: a regulatory system in yeast. Microbiology, 44(2), pp.149–156.

11. Derrien, D., Plain, C., Courty, P.E., Gelhaye, L., Moerdijk-Poortvliet, T.C., Thomas, F., Versini, A., Zeller, B., Koutika, L.S., Boschker, H.T. and Epron, D., 2014. Does the addition of labile substrate destabilise old soil organic matter?. Soil Biology and Biochemistry, 76, pp.149–160.

12. Dijkstra, P., Salpas, E., Fairbanks, D., Miller, E.B., Hagerty, S.B., van Groenigen, K.J., Hungate, B.A., Marks, J.C., Koch, G.W. and Schwartz, E., 2015. High carbon use efficiency in soil microbial communities is related to balanced growth, not storage compound synthesis. Soil Biology and Biochemistry, 89, pp.35–43.

13. Dilly, O. and Zyakun, A., 2008. Priming effect and respiratory quotient in a forest soil amended with glucose. Geomicrobiology Journal, 25(7-8), pp.425–431.

14. Edwards, V.H., 1970. The influence of high substrate concentrations on microbial kinetics. Biotechnology and Bioengineering, 12(5), pp.679–712.

15. Frey, S.D., Lee, J., Melillo, J.M. and Six, J., 2013. The temperature response of soil microbial efficiency and its feedback to climate. Nature Climate Change, 3(4), pp.395–398.

16. Geyer, K.M., Dijkstra, P., Sinsabaugh, R. and Frey, S.D., 2019. Clarifying the interpretation of carbon use efficiency in soil through methods comparison. Soil Biology and Biochemistry, 128, pp.79–88.

17. Hagerty, S.B., Van Groenigen, K.J., Allison, S.D., Hungate, B.A., Schwartz, E., Koch, G.W., Kolka, R.K. and Dijkstra, P., 2014. Accelerated microbial turnover but constant growth efficiency with warming in soil. Nature Climate Change, 4(10), pp.903–906.

18. Han, K. and Levenspiel, O., 1988. Extended Monod kinetics for substrate, product, and cell inhibition. Biotechnology and bioengineering, 32(4), pp.430–447.

19. Hobbie, J.E. and Hobbie, E.A., 2013. Microbes in nature are limited by carbon and energy: the starving-survival lifestyle in soil and consequences for estimating microbial rates. Frontiers in microbiology, 4, p.324.

20. Iooss, B., Da Veiga, S., Janon, A., Pujol, G., et al., 2020. sensitivity: Global Sensitivity Analysis of Model Outputs. R package version 1.20.0. https://CRAN.R-project.org/package=sensitivity

21. Johnson, C.K., Vigil, M.F., Doxtader, K.G. and Beard, W.E., 1996. Measuring bacterial and fungal substrate-induced respiration in dry soils. Soil Biology and Biochemistry, 28(4-5), pp.427–432.

22. Kempes, C.P., van Bodegom, P.M., Wolpert, D., Libby, E., Amend, J. and Hoehler, T., 2017. Drivers of bacterial maintenance and minimal energy requirements. Frontiers in microbiology, 8, p.31.

23. Kreft, J.U. and Bonhoeffer, S., 2005. The evolution of groups of cooperating bacteria and the growth rate versus yield trade-off. Microbiology, 151(3), pp.637–641.

24. Lennon, J.T. and Cottingham, K.L., 2008. Microbial productivity in variable resource environments. Ecology, 89(4), pp.1001–1014.

25. Lipson, D.A., 2015. The complex relationship between microbial growth rate and yield and its implications for ecosystem processes. Frontiers in microbiology, 6, p.615.

26. Lin, Q. and Brookes, P.C., 1999. An evaluation of the substrate-induced respiration method. Soil Biology and Biochemistry, 31(14), pp.1969–1983.

27. Manzoni, S., Taylor, P., Richter, A., Porporato, A. and Ågren, G.I., 2012. Environmental and stoichiometric controls on microbial carbon-use efficiency in soils. New Phytologist, 196(1), pp.79–91.

28. Molenaar, D., Van Berlo, R., De Ridder, D. and Teusink, B., 2009. Shifts in growth strategies reflect tradeoffs in cellular economics. Molecular systems biology, 5(1), p.323.

29. Nakamoto, T. and Wakahara, S., 2004. Development of Substrate Induced Respiration (SIR) Method Combined with Selective Inhibition for Estimating Fungal andBacterial Biomass in Humic Andosols. Plant production science, 7(1), pp.70–76.

30. Olsen, S. and Sommers, L.E., 1982. Phosphorus. Methods of soil analysis, pp.403–430.

31. Panikov, N.S. and Sizova, M.V., 1996. A kinetic method for estimating the biomass of microbial functional groups in soil. Journal of Microbiological Methods, 24(3), pp.219–230.

32. Pfeiffer, T., Schuster, S. and Bonhoeffer, S., 2001. Cooperation and competition in the evolution of ATP-producing pathways. Science, 292(5516), pp.504–507.

33. Reischke, S., Kumar, M.G. and Bååth, E., 2015. Threshold concentration of glucose for bacterial growth in soil. Soil Biology and Biochemistry, 80, pp.218–223.

34. Schneckenberger, K., Demin, D., Stahr, K. and Kuzyakov, Y., 2008. Microbial utilization and mineralization of [14C] glucose added in six orders of concentration to soil. Soil Biology and Biochemistry, 40(8), pp.1981–1988.

35. Schnecker, J., Bowles, T., Hobbie, E.A., Smith, R.G. and Grandy, A.S., 2019. Substrate quality and concentration control decomposition and microbial strategies in a model soil system. Biogeochemistry, 144(1), pp.47–59.

36. Schumacher, B.A., 2002. Methods for the determination of total organic carbon (TOC) in soils and sediments.

37. Schuster, S., Boley, D., Möller, P., Stark, H. and Kaleta, C., 2015. Mathematical models for explaining the Warburg effect: a review focussed on ATP and biomass production. Biochemical Society Transactions, 43(6), pp.1187–1194.

38. Sinsabaugh, R.L., Turner, B.L., Talbot, J.M., Waring, B.G., Powers, J.S., Kuske, C.R., Moorhead, D.L. and Follstad Shah, J.J., 2016. Stoichiometry of microbial carbon use efficiency in soils. Ecological Monographs, 86(2), pp.172–189.

39. Steinweg, J.M., Plante, A.F., Conant, R.T., Paul, E.A. and Tanaka, D.L., 2008. Patterns of substrate utilization during long-term incubations at different temperatures. Soil biology and biochemistry, 40(11), pp.2722–2728.

40. Sterner, R.W. and Elser, J.J., 2002. Ecological stoichiometry: the biology of elements from molecules to the biosphere. Princeton university press.

41. Swain, A. and Fagan, W.F., 2018. A mathematical model of the Warburg Effect: Effects of cell size, shape and substrate availability on growth and metabolism in bacteria. Math. Biosci. Eng, 16, pp.168–186.

42. Taylor, M.D., 2000. Determination of total phosphorus in soil using simple Kjeldahl digestion. Communications in Soil Science and Plant Analysis, 31(15-16), pp.2665–2670.

43. Tian, J., Pausch, J., Yu, G., Blagodatskaya, E., Gao, Y. and Kuzyakov, Y., 2015. Aggregate size and their disruption affect 14C-labeled glucose mineralization and priming effect. Applied Soil Ecology, 90, pp.1–10.

44. Vander Heiden, M.G., Cantley, L.C. and Thompson, C.B., 2009. Understanding the Warburg effect: the metabolic requirements of cell proliferation. science, 324(5930), pp.1029–1033.

45. Voet, D., 2004; Biochemistry. Hoboken.

46. Wilke, B.M., 2005. Determination of chemical and physical soil properties. In Monitoring and assessing soil bioremediation (pp. 47–95). Springer, Berlin, Heidelberg.

47. Young, K.D., 2006. The selective value of bacterial shape. Microbiology and molecular biology reviews, 70(3), pp.660–703.

48. Zhuang, K., Vemuri, G.N. and Mahadevan, R., 2011. Economics of membrane occupancy and respiro-fermentation. Molecular systems biology, 7(1), p.500.

